# Cigarette Smoke and Decreased DNA Repair by Xeroderma Pigmentosum Group C Use a Double Hit Mechanism for Epithelial Cell Lung Carcinogenesis

**DOI:** 10.1101/2025.02.22.639660

**Authors:** Nawar Al Nasrallah, Bowa Lee, Benjamin M. Wiese, Marie N. Karam, Elizabeth A. Mickler, Huaxin Zhou, Nicki Paolelli, Robert S. Stearman, Mark W. Geraci, Catherine R. Sears

## Abstract

Emerging evidence suggests a complex interplay of environmental and genetic factors in non-small cell lung cancer (NSCLC) development. Among these factors, compromised DNA repair plays a critical but incompletely understood role in lung tumorigenesis and concurrent lung diseases, such as chronic obstructive lung disease (COPD).

In this study, we investigated the interplay between cigarette smoke, DNA damage and repair, focusing on the Nucleotide Excision Repair (NER) protein Xeroderma Pigmentosum Group C (XPC). We found decreased XPC mRNA expression in most NSCLCs compared to subject-matched, non-cancerous lung. In non-cancerous bronchial epithelial cells, cigarette smoke decreased NER, increased total DNA damage and resultant apoptosis, each exacerbated by XPC deficiency. In contrast, lung cancer cells exhibit greater resilience to cigarette smoke, requiring higher doses to induce comparable DNA damage and apoptosis, and are less reliant on XPC expression for survival. Importantly, XPC protects against chromosomal instability in benign bronchial epithelial cells, but not in lung cancer cells. Our findings support a “double hit” mechanism wherein early decreased XPC expression and resultant aberrant DNA repair, when combined with cigarette smoke exposure, may lead to loss of non-malignant epithelial cells (as observed in COPD), and contributes to early NSCLC transition through altered DNA damage response.

## Introduction

Lung cancer remains the leading cause of cancer-related fatalities in the United States and worldwide, with cigarette smoking the most well-established risk factor for lung cancer^1^. Cigarette smoke contains a complex mix of carcinogens, including polycyclic aromatic hydrocarbons and nitrosamines, that directly damage cellular DNA^2^. This DNA damage manifests as adducts, strand breaks, and oxidative lesions, constantly challenging the genomic integrity of lung epithelial cells^3^. DNA repair pathways are critical for repairing such damage, but aberrations can lead to unrepaired DNA damage, replication errors, genotoxic stress, and ultimately cancer ^4^. Decreased DNA repair can result in differential DNA damage responses, including altered apoptosis, autophagy, and/or senescence^5^. Additionally, a shift from classic to low-fidelity DNA repair pathways may further augment genome instability, ultimately contributing to both lung cancer and COPD/emphysema^6^.

Several DNA repair pathways are implicated in cancer, including nucleotide excision repair (NER), which repairs bulky DNA intra-strand adducts from cigarette smoke, and base excision repair (BER), which repairs nucleotide damage from oxidation, deamination and alkylation ^7^.

Xeroderma Pigmentosum Group C (XPC) is a crucial protein in recognizing DNA strand distortion and initiating global genomic nucleotide excision repair (GG-NER), a process critical to genome-wide repair of bulky DNA lesions such as those caused by cigarette smoke ^5,8^. It also plays a role in base excision repair (BER) and protects against carcinogen-induced oxidative DNA damage in mice^5,9^. We found that XPC protects against carcinogen-induced lung adenocarcinoma in mice^8,10^. Additionally, XPC deficiency in mice exposed to the carcinogen N-nitroso-tris-chloroethylurea (NTCU) is associated with an increased frequency and size of lung squamous cell carcinomas and earlier progression of pre-malignant squamous dysplasia^11^. Chronic (6 months) cigarette smoke exposure through a smoking chamber decreases *Xpc* gene expression in mice and XPC deficiency increases emphysema-like lung disease in mice with age, exacerbated by cigarette smoke exposure ^5,6,12^. Understanding these mechanisms is vital for identifying preventive and therapeutic targets to combat cigarette smoke-associated lung carcinogenesis.

Our central hypothesis posits that decreased XPC mRNA expression in lung cancer cells leads to reduced DNA repair, increased DNA damage, and genomic instability, which ultimately contribute to lung cancer development. This paper explores the impact of cigarette smoke on XPC-mediated DNA damage and repair in bronchial epithelial and NSCLC cell lines, revealing a novel differential impact on pre-malignant bronchial epithelial and NSCLC cells. Additionally, we identify a role of XPC in maintenance of genomic stability in benign bronchial epithelial cells but not NSCLC cells. This study provides insights into the potential mechanistic role of XPC-mediated DNA repair in the development of both non-small cell lung cancer and emphysema.

## Results

### 1. XPC deficiency differentially impacts cancer and bronchial epithelial cell susceptibility to cell death through apoptosis during cigarette smoke exposure

We previously observed both accelerated emphysema and NSCLC development in XPC-deficient (XPC KO) mice exposed to cigarette smoke (CS) and CS-carcinogens compared to those with wild-type XPC expression (XPC WT)^11-13^. We found that XPC protected against CS-induced apoptotic cell death in non-cancerous bronchial epithelial cells (Beas-2B) and in chronic CS-exposed mouse lung with emphysema-like changes^12^. Additionally, Beas-2B cells modified by stable lentiviral knock-down of XPC (shXPC), have decreased clonogenic survival to CS extract (CSE) driven by increased apoptosis when compared to non-targeted, scramble controls (shCtrl) Beas-2B cells^12^. However, cancer cells are characterized by inhibition of apoptosis, raising the question of whether XPC differentially impacts cellular response to CS in benign epithelial and cancer cells.

We evaluated the impact of XPC on survival of NSCLC cell lines to CS. Non-small cell lung cancer cell lines (A549, H1299 and H520) were modified by stable XPC knock-down (shXPC) or non-targeted control (shCtrl) as published^12^. Clones used for these experiments had XPC mRNA expression decreased by 55-64% by RT-qPCR and confirmatory Western blot analysis (Supplemental Figure 1). Compared to Beas-2B cells, clonogenic survival was increased in all NSCLC cell lines by titratable CSE concentrations, with an inverse dose-response survival association (Figure 1 and previously published^12^). Unlike Beas-2B cells, in which shXPC was associated with decreased clonogenic survival, in NSCLC cell lines (A549, H1299 and H520), shXPC led to no statistically significant difference in clonogenic survival compared to shCtrl (Figure 1). We further confirmed the mechanism for this response by measuring apoptotic response to CSE by Annexin V-PI staining and expression of the activated (cleaved) form of apoptosis factors caspase 9, caspase 3 and PARP (Figure 2). In H1299 cells, apoptosis was increased by exposure to CSE, but was not impacted by shXPC compared to shCtrl (Figure 2 A-C). H520 cells showed an increase in apoptosis with exposure to CSE, however, XPC knock-down did not alter apoptosis with high (12.5%) CSE exposure, a level associated with 13.5-19.2% clonogenic survival in these cell lines (Figure 2 D-F). Only exposure to high (20%) CSE, a concentration associated with almost no survival, is associated with increases in apoptosis as measured by Annexin V (Figure 2 D), cleaved caspase 3 and cleaved PARP in shXPC (Figure 2F). A549 cells showed CSE-induced apoptosis increased in shCtrl but not in shXPC cells, with decreased activation of the apoptotic pathway in shXPC cells (Figure 2 G-I). Together, these findings suggest that XPC differentially impacts cellular response to CSE in NSCLC compared to benign bronchial epithelial cells, leading to differences in cell survival and apoptosis.

**Figure 1:**
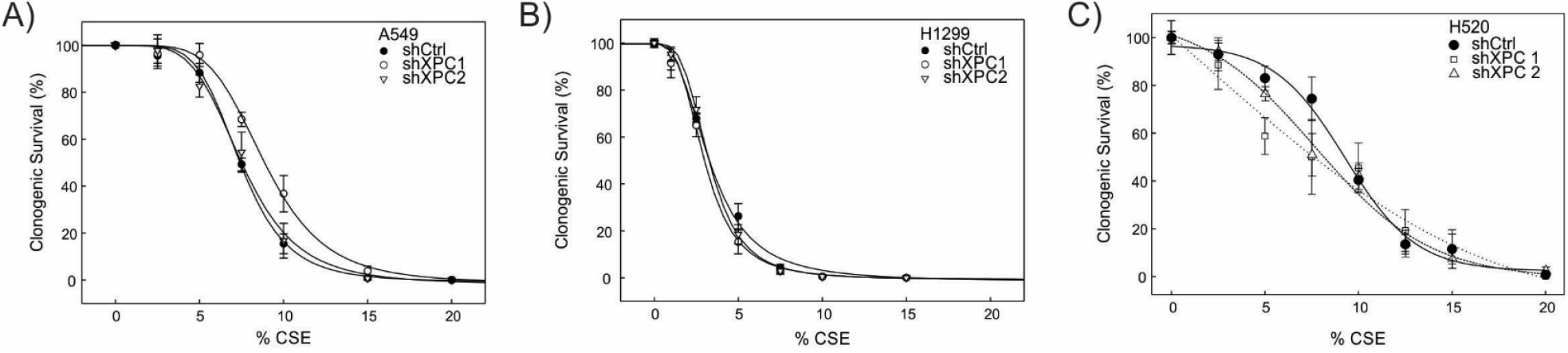
Clonogenic Survival in Response to Increasing Concentrations of CS Extract (CSE) Clonogenic survival assays were conducted on XPC knockdown (shXPC) cells in comparison to the control (shCtrl) under varying concentrations of CS extract (CSE) for A) A549, B) H1299 and C) H520 non-small cell lung cancer (NSCLC) cell lines. The survival data is presented as individual points with error bars indicating standard error of the mean (±SEM) and fitted to a best-fit four-parameter logistic curve. Statistical differences in CSE survival were assessed for each shXPC cell line compared to shCtrl using a two-way ANOVA with repeated measures with no significant differences were observed in the clonogenic survival of the human NSCLC cell lines.

**Figure 2:**
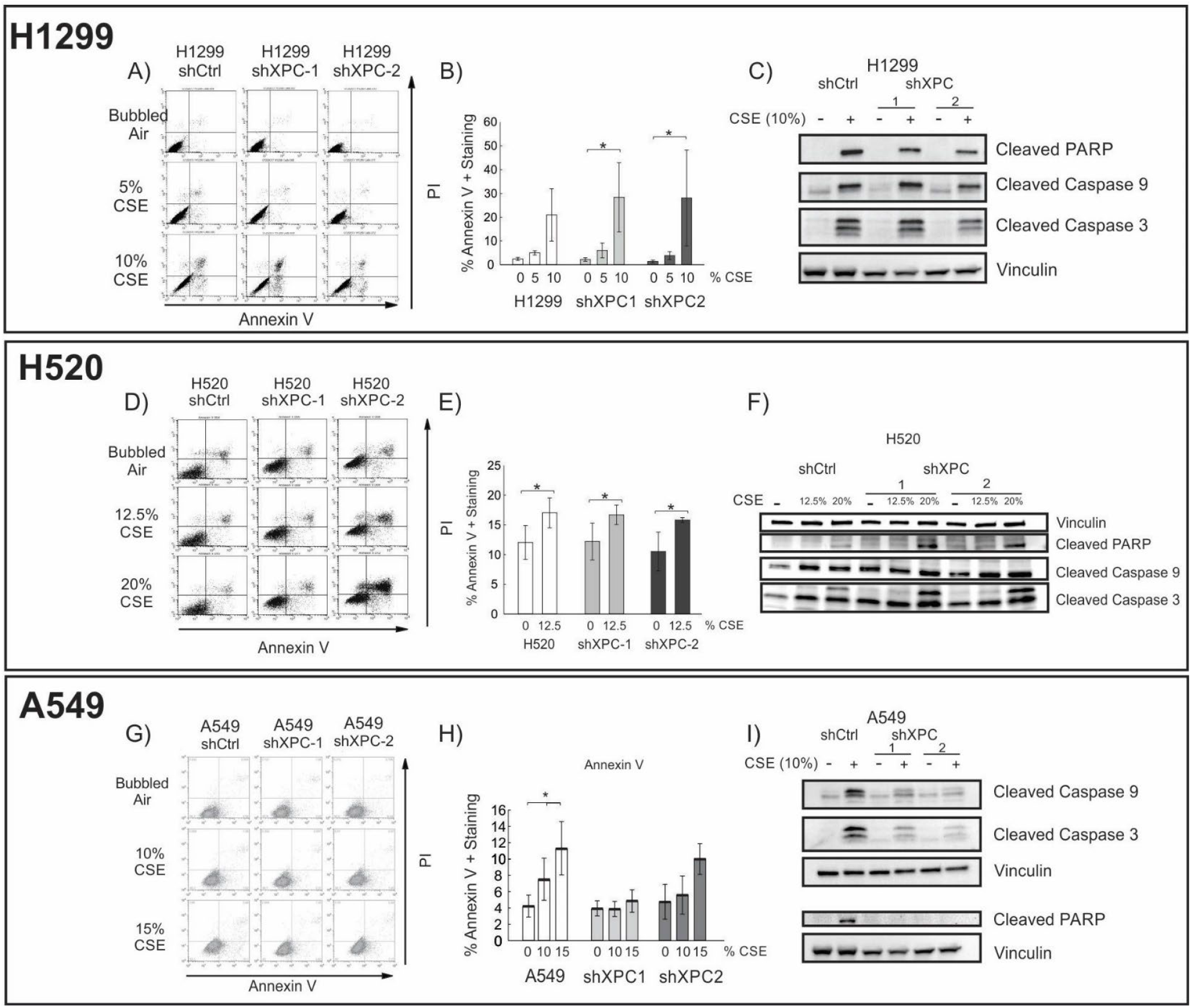
XPC Impact to Cigarette Smoke Extract – Induced Apoptosis in NSCLC Cell Lines. Flow Cytometry showing Annexin V-PI stain and quantification of early and late apoptotic cells in H1299 (A, B) and H520 (D, E) and A549 (G, H) cell lines. Confirmation of apoptosis by Western blot for activated PARP, Caspase 9, and Caspase 3 in C) H1299, F) H520 and I) A549 NSCLC cell lines.

### 2. XPC protects against cigarette smoke induced total and oxidative DNA damage in benign and cancerous lung cells

Both NER and BER functions of XPC are being implicated in protecting against lung tumorigenesis^10,13^. We therefore sought to determine the impact of XPC on CS-induced DNA damage, and whether this differed in benign bronchial epithelial compared to NSCLC cells. Using our shXPC and shCtrl modified bronchial epithelial (Beas-2B) and NSCLC cell lines (A549, H1299, H520), we measured levels of DNA damage to cigarette smoke extract (CSE) or air control (AC). Regardless of XPC expression, all NSCLC cell lines required treatment with higher concentrations of CSE to develop similar levels of DNA damage by alkaline Comet assay compared to benign bronchial epithelial cells (Beas-2B) (Figure 3). DNA damage increased with higher concentrations of CSE and with XPC knock-down in all cell lines (Figure 3). We investigated whether there was a difference in CSE-induced oxidative DNA damage using hOGG1 FLARE Comet assay that specifically-measures 8-oxo-7,8-dihydro-2’-deoxyguanosine (8-oxoG). As with total DNA damage, we found that Beas-2B cells were more susceptible to CSE-induced oxidative DNA damage compared to NSCLC cell lines. XPC protected against 8-oxoG DNA damage in both Beas-2B and NSCLC (H1299 and A549) cell lines (Figure 4). These findings suggest that XPC plays a protective role against oxidative and total DNA damage in both NSCLC and benign bronchial epithelial cells.

**Figure 3:**
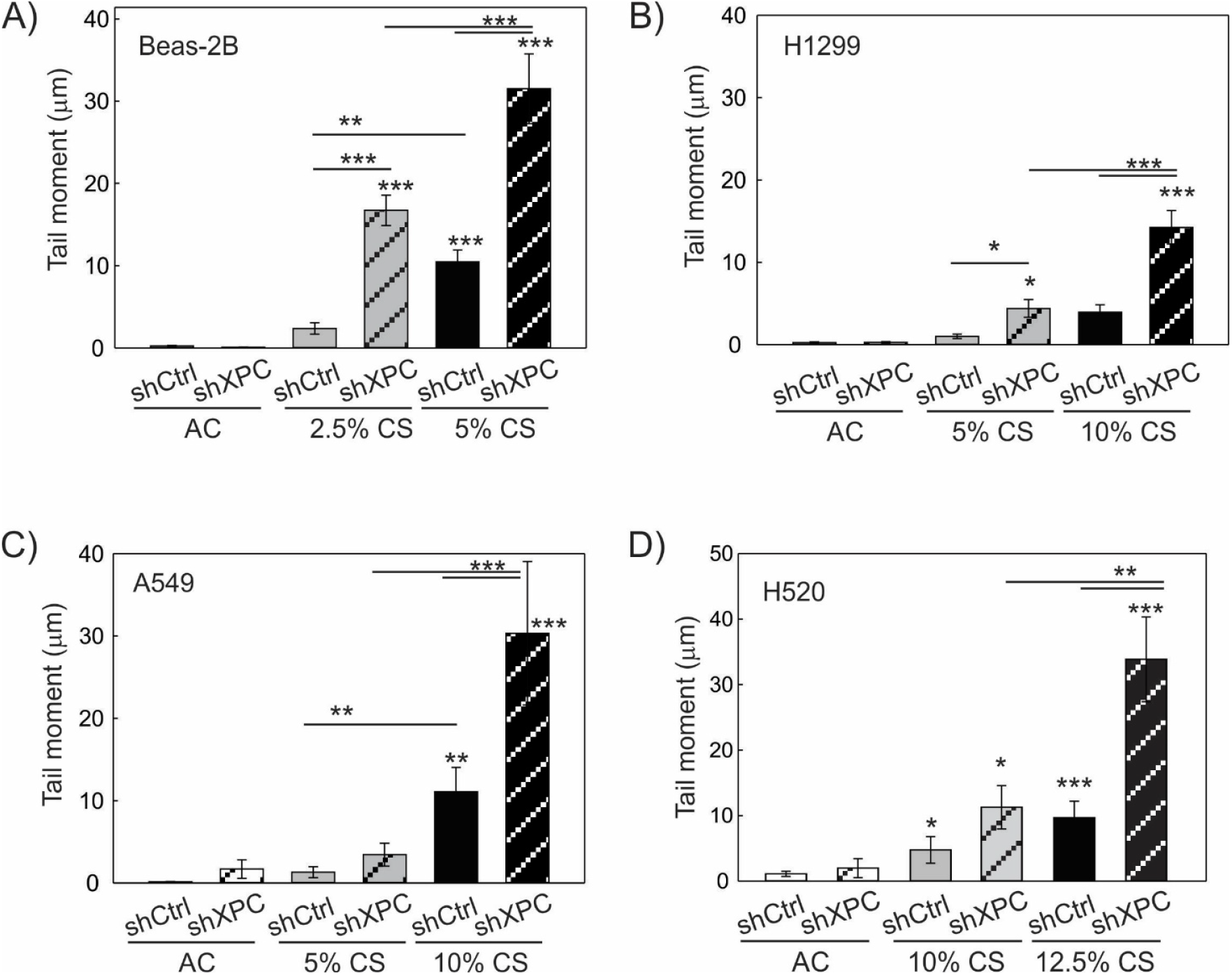
Total DNA Damage Assessed by Alkaline Comet Assay. Total DNA damage measured following 24 hours of exposure to cigarette smoke extract (CS) or filtered air (AC) in A) benign epithelial cells (Beas-2B), B) A549 lung adenocarcinoma cell line, C) H1299 lung adenocarcinoma cell line and D) H520 lung adenocarcinoma cell line modified by XPC knock-down (shXPC) or scramble control (shCtrl). Note increased DNA damage correlates to increasing CS concentrations and is further increased by shXPC compared to shCtrl in all cell lines. Mean +/-SD from 3 separate experiments, *p<0.05, **p<0.01, ***p<0.001 by 2-way ANOVA.

**Figure 4:**
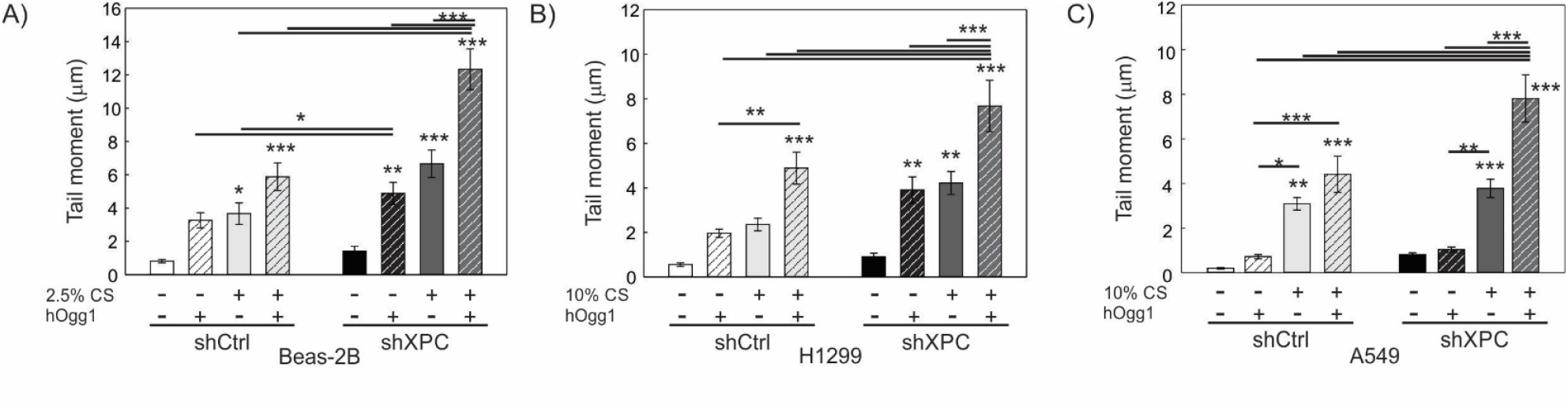
Effect of XPC in Oxidative DNA Damage in Benign and Cancerous Lung Epithelial Cells. Analysis of 8-hydroxy-2’-deoxyguanosine (8-OHdG) adducts following 24-hour exposure to cigarette smoke extract (CS, +) or filtered air (−) using FLARE Comet Assay and human 8-hydroxyguanine DNA-glycosylase 1 (hOGG1) ^45^ in A) Benign bronchial epithelial cells (Beas-2B) modified by XPC knock-down (shXPC) or scramble control (shCtrl), B) H1299 NSCLC cells modified by shXPC compared to shCtrl and C) A549 NSCLC cells modified by shXPC compared to shCtrl. Note the amount of oxidative DNA damage increases with CS concentration and exhibits a more pronounced impact in shXPC compared to shCtrl in all cell lines. Mean +/-SD from 3 independent experiments, *p<0.05, **p<0.01, ***p<0.001 by 2-way ANOVA.

### 3. Micronucleus formation increases with XPC knockdown in Beas-2B but not malignant NSCLC cell lines

Micronuclei are small, DNA-containing membrane-bound structures separated from the cell nucleus containing chromosomes or chromosome fragments unable to be incorporated into the nucleus during mitosis. These can be caused by DNA strand breaks directly caused by DNA damage or developed when damage impairs DNA replication, and are associated with genomic instability, mutagenesis and cancer development ^14^. Chromosomal aberrancies are structural and numerical changes in chromosomes, often caused by misrepair of double strand DNA breaks, leading to instability that drives cancer development and growth ^15^. A cytokinesis-block micronucleus (CBMN) assay was completed to assess the impact of XPC on genomic instability in human bronchial epithelial (Beas-2B) and NSCLC cells. Both micronuclei and nuclear aberrancies increased significantly by exposure to CSE, as compared by 2-way ANOVA (p=0.003, micronuclei; p=0.002, nuclear aberrancies) (Figure 5 A-C). The percent cells with micronuclei were significantly increased by XPC knockdown in Beas-2B cells (Figure 5B, p<0.001), but no XPC effect was observed on non-specific nuclear aberrancies (Figure 5C, p=0.181).

**Figure 5:**
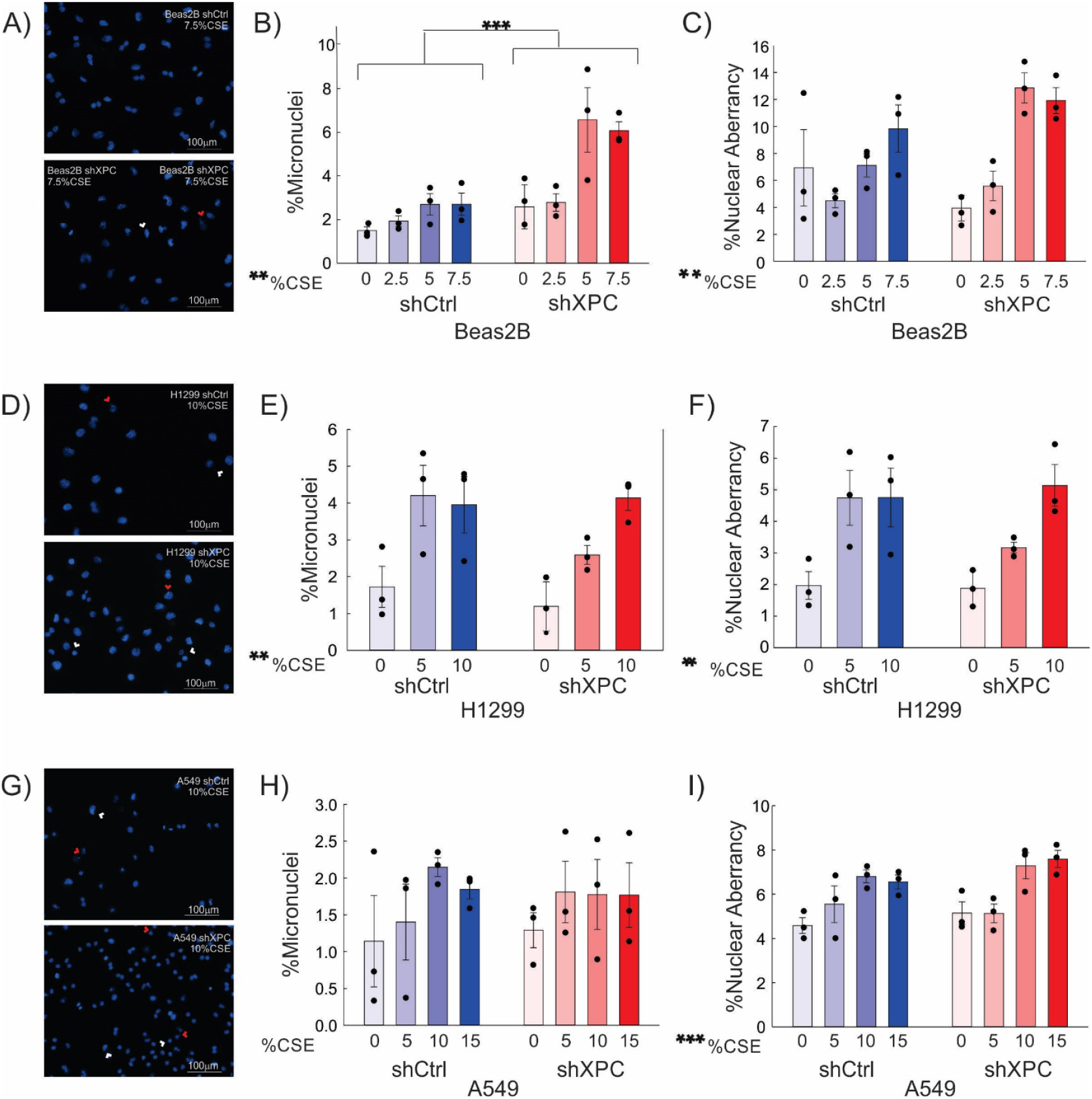
Micronuclei and Nuclear Aberrancies. A) Representative DAPI images of nuclear aberrancy (nuclear blebs or bridges, red arrow) and micronuclei (white arrow) in Beas-2B cells exposed to CSE. B and C) Quantification of % Beas-2B cells with micronuclei (B) and % cells with nuclear aberrancies (C). Results are also shown for H1299 (D, E, F) and A549 (G, H, I). shXPC = lentiviral XPC knock-down. shCtrl = scrambled shRNA control. **= p<0.01; ***=p<0.001 by 2-way ANOVA.

In both H1299 and A549 NSCLC cell lines, nuclear aberrancies were augmented by CSE by two-way ANOVA (Figure 5 F and I, p=0.004, H1299; p<0.001, A549). Micronuclei formation increased with CSE treatment in H1299 cells (p=0.005) (Figure 5 E); this effect was not observed in A549 cells (p=0.325) (Figure 4H).

Knockdown of XPC had no effect on micronuclei formation in either H1299 or A549 cell lines exposed to increasing concentrations of CSE (Figure 5 E and H). Our findings suggest that XPC exhibits a protective role against development of development of mutagenic genome alterations in benign bronchial epithelial cells, but its role in protection against development of genomic instability is diminished in malignant cells.

### 4. Cigarette smoke decreases DNA repair in benign bronchial but not malignant lung epithelial cells

To further investigate the differential impact of CS in benign and malignant lung epithelial cells, we next investigated DNA repair capacity in response to CS exposure. We measured percent nucleotide excision repair (NER) using a host-cell reactivation assay. NER efficiency was determined by transfection of a UV-modified, green fluorescent protein (GFP)-expressing plasmid requiring NER repair of UV lesions to express GFP compared to GFP expression from unmodified plasmid transfection. Cells were co-transfected with an unmodified, E2-Crimson expressing plasmid (transfection control) and percent NER efficiency determined by the ratio of GFP to E2-Crimson in cells by flow-cytometry^16^. CSE exposure for 24 hours was associated with decreased NER in benign Beas-2B cells (Figure 6A). In contrast, CSE exposure did not significantly affect NER in the A549 NSCLC cell line, although A549 cells appeared to have a lower DNA repair at baseline (Figure 6B). We hypothesized that the effect of CSE on DNA repair may be due to decreased XPC expression, as previously observed in the lungs of C57Bl/6 mice exposed to 6 months of cigarette smoke^17^. We treated Beas-2B cells with increasing concentrations of CSE for 24 hours and found that XPC mRNA expression was decreased by RT-qPCR (Figure 6C). In conclusion, the differential effect of CS on benign cells, leading to decreased DNA repair, could explain its role in the progression towards malignant cells, which exhibit inherently lower DNA repair capabilities.

**Figure 6:**
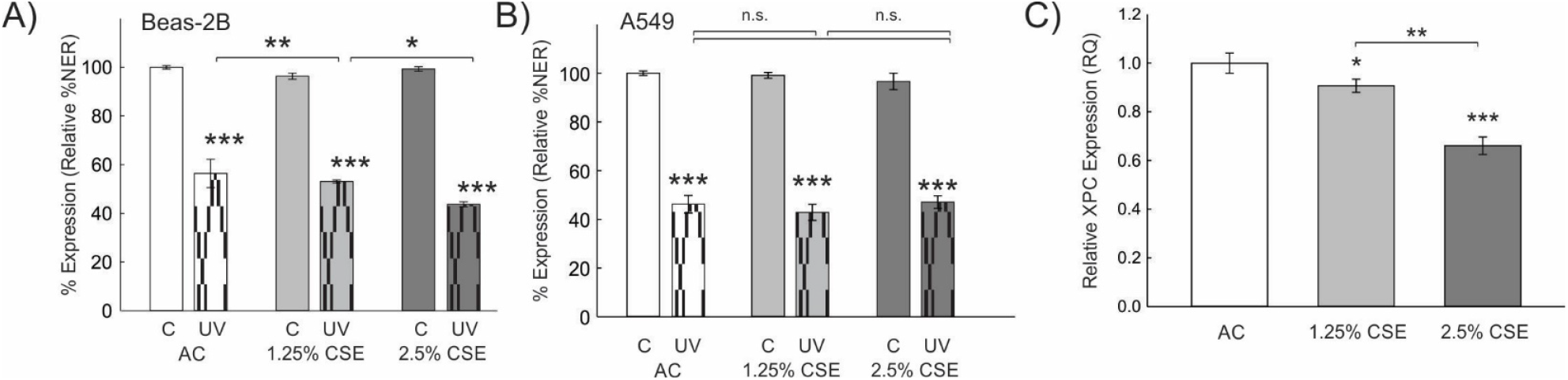
Effect of Cigarette Smoke Extract (CSE) on Nucleotide Excision Repair (%NER) and XPC gene Expression. Relative NER efficiency of UV-modified plasmid (pMAX-GFP, UV) or unmodified plasmid (C) in cells treated with increasing concentrations of CSE or air control (AC). A) %NER in bronchial epithelial cells (Beas-2B) is higher at baseline and decreases significantly after treatment with CSE. B) %NER in human A549 NSCLC cell line is decreased at baseline and not significantly altered by CSE. Shown as mean +/-standard deviation from 3 separate experiments. UV = ultraviolet. *p<0.05, **p<0.01, ***p<0.001 by 2-Way ANOVA. C) XPC mRNA expression by RT-qPCR is decreased in Beas-2B cells exposed to CSE for 24 hours. Statistical analysis by one-way ANOVA using dCt values.

### 5. XPC gene expression is decreased in human lung adenocarcinoma compared to non-cancerous lung

Given the above findings, we hypothesized that XPC gene expression would be decreased in human NSCLCs adenocarcinomas compared to non-cancerous lung specimens. We first analyzed this in lung adenocarcinomas. Gene expression data from the TCGA database was used to determine XPC mRNA expression in 502 lung adenocarcinoma samples and 59 benign lung samples. We found a significant decrease in XPC gene expression in lung adenocarcinomas compared to unmatched benign lung samples (Figure 7A). To account for cigarette smoking as a possible cause of decreased XPC mRNA expression^8^, we also evaluated XPC gene expression based on cigarette smoking status associated with each of these TCGA samples. Although the number of normal lung tissue was low for current and never cigarette smoking, XPC gene expression was significantly decreased in lung adenocarcinomas compared to normal lung irrespective of cigarette smoking status (Supplemental Figure 2) and gender (data not shown).

**Figure 7.**
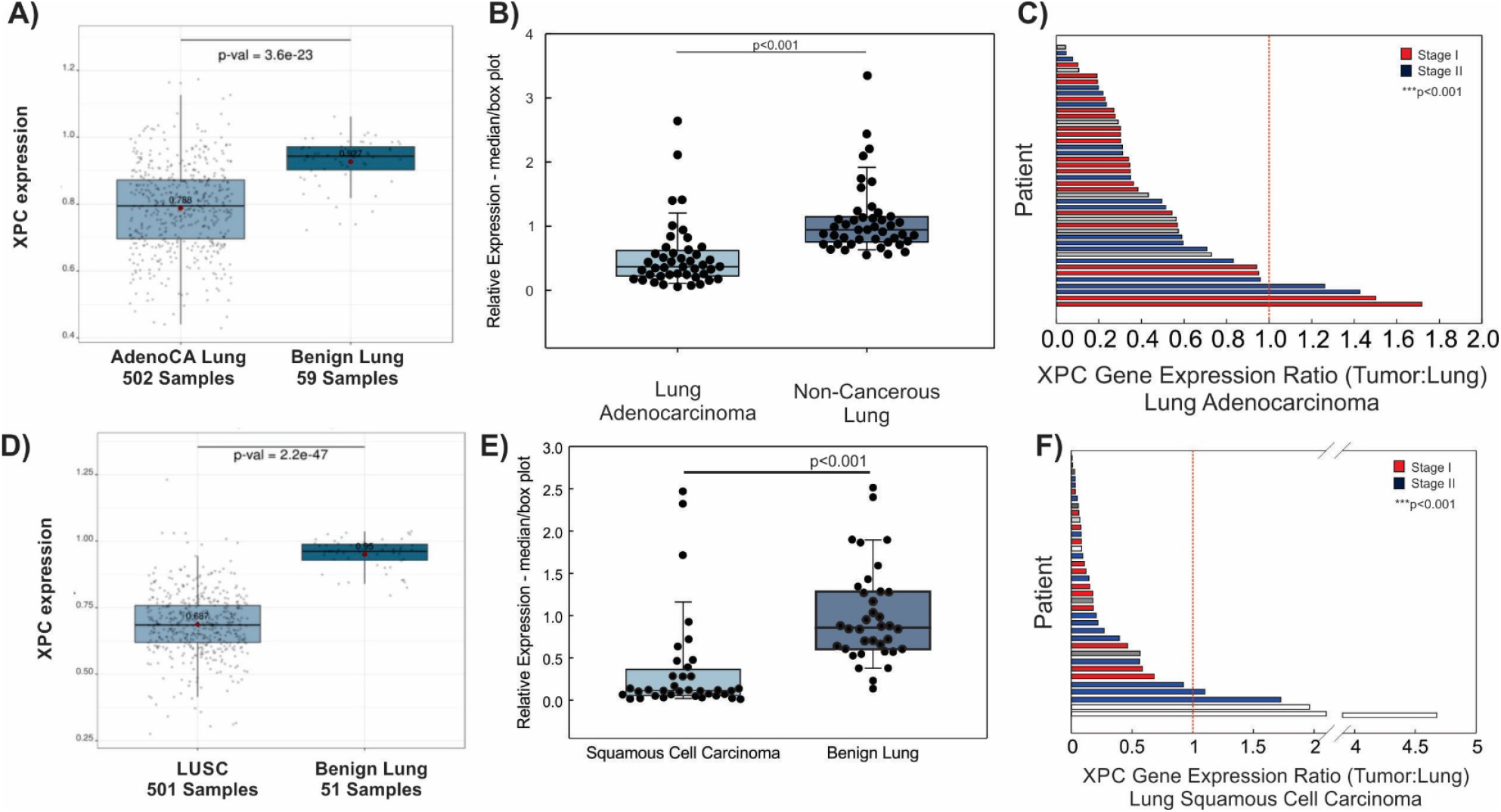
XPC Gene Expression is Decreased in Human Non-Small Cell Lung Cancers Compared to Normal Subject-Matched Lung. A) XPC mRNA expression decreased in unmatched samples from lung adenocarcinoma (AdenoCA) compared to benign lung from the TCGA database. B) Decreased XPC mRNA expression in frozen lung adenocarcinoma compared to non-cancerous (benign) lung. Box plot with median and 25-75%, whiskers at 10% and 95%. C) Ratio of XPC mRNA expression in lung adenocarcinoma to subject matched benign lung resected at the time of surgery, individual subjects shown on Y-axis. D) XPC mRNA expression decreased in unmatched samples from the TCGA database of lung squamous cell carcinoma (LUSC) and benign lung. E) Decreased median XPC mRNA expression in frozen lung squamous cell carcinoma compared to benign lung. Box plot with median and 25-75%, whiskers at 10% and 95%. F) Ratio of XPC mRNA expression in lung squamous cell carcinoma to subject-matched benign lung resected at the time of surgery, individual subjects shown on Y-axis. ***p<0.001 by one way ANOVA.

Oncogenic mutations and copy number variations have been detected in non-cancerous lung tissue and epithelial brush/biopsy samples from subjects with lung cancer (known as the “field effect “)^18-20^. To account for this “field effect” and to reduce the potential bias introduced by evaluating gene expression across different individuals, we determined XPC mRNA expression in frozen human lung adenocarcinoma compared to subject-matched non-cancerous adjacent lung from 44 subjects. Supplemental Table 1 shows clinical characteristics. Median XPC mRNA expression was decreased in human lung adenocarcinoma compared to non-cancerous, resected lung from the same individual (Figure 7 B and C). The ratio of XPC mRNA expression was decreased in lung adenocarcinoma tissue compared to subject-matched benign lung irrespective of cigarette smoking status and stage at the time of diagnosis (Supplemental Figure 2 and Figure 7C).

### 6. XPC is decreased in human lung squamous cell carcinoma compared to non-cancerous lung

Little is known about the role of XPC in lung squamous cell carcinoma (LUSC) development. In a mouse model of chronic CS exposure, we observed that pre-cancerous squamous dysplastic changes preceded lung cancer development in XPC-deficient mice exposed to chronic CS, and that mice deficient in XPC display accelerated progression of premalignant squamous dysplasia, associated with earlier development, larger size and higher incidence^11,12^. We therefore hypothesized that XPC mRNA would be decreased in lung squamous cell carcinoma compared to normal lung tissue similar to that observed in lung adenocarcinoma. Using archived gene expression data from 501 squamous cell carcinomas and 51 non-cancerous lung specimens stored in the TCGA database, we observed decreased XPC mRNA expression in lung squamous cell carcinoma compared to normal lung samples (Figure 7D). Decreased XPC mRNA was observed in both male and female genders (not shown). Lung squamous cell carcinomas occur almost exclusively in those with a cigarette smoking history; both current and former cigarette smoking was associated with decreased XPC gene expression in squamous cell carcinoma compared to normal lung tissue (Supplemental Figure 2).

We again wanted to control for potential biases associated with inter-individual gene expression comparisons and possible “field effect” of cigarette smoking in lung squamous cell carcinoma samples. For this reason, we evaluated XPC mRNA expression by RT-qPCR in frozen lung squamous cell carcinoma and matched resected lung specimens from 36 subjects. Included tumor samples represented Stages I-III squamous cell carcinoma. As expected, most subjects were current or former smokers of cigarettes (Supplemental Table 2). Overall, median XPC mRNA expression was decreased compared to non-cancerous lung, recapitulating data from the TCGA database (Figure 7E). Importantly, lung squamous cell carcinomas had decreased XPC gene expression ratio compared to subject-matched adjacent, non-cancerous lung (Figure 7F), further confirming that observed decreases in XPC in NSCLCs is tumor-related rather than due to inter-individual changes.

## Discussion

In this study, we investigated the impact of XPC on cigarette smoke-induced cellular changes in benign pulmonary epithelial and NSCLC cells. Our key findings highlight that cigarette smoke affects benign bronchial cells differently compared to malignant cell lines, evidenced by greater DNA damage in benign cells versus malignant lines, both in terms of oxidative and overall DNA damage. XPC knock-down and cigarette smoke, known contributors to genomic instability, were both shown to specifically augment cigarette smoke-induced chromosomal breaks, manifested as micronuclei and chromosomal aberrancy in pre-malignant Beas-2B bronchial epithelial cells^12,21-23^. These results are pivotal to understanding carcinogenesis mechanisms in lung cancer, in which a growing body of evidence links XPC polymorphisms to cancer risk^24-26^. This may also inform carcinogenesis mechanisms across other cancer types, as an increased prevalence of XPC deletions or polymorphisms has been described in lung, prostate, bladder, hematologic, and other cancers^5^. Decreased XPC expression may result in a shift from high-fidelity DNA repair mechanisms to low-fidelity ones, driving genomic instability, as suggested by the finding of increased micronuclei in XPC knock-down Beas-2B cells exposed to cigarette smoke extract.

Moreover, we found that CSE decreased NER in benign bronchial cells due to reduced XPC protein expression. This is consistent with the findings of others who have demonstrated decreased NER capacity following cigarette smoke exposure^27^. Our study suggests that cigarette smoke exposure leads to decreased XPC mRNA expression, exacerbates total and oxidative DNA damage, hinders NER, and may contribute to lung cancer development. The precise mechanism of genomic instability remains challenging to pinpoint due to XPC’s multifaceted roles beyond traditional NER, including oxidative damage repair through base excision repair (BER), interaction with p53 and other potential mechanisms^5^. Interestingly, Lindbergh et al. found that nuclear blebs were more likely to form from interstitial DNA, whereas micronuclei occurred when the terminal segment of a chromosome broke off^28^. Future research should focus on the mechanistic role of XPC in micronucleus formation and genomic instability, and potentially novel functions of XPC in replication stress.

In contrast, NSCLC cell lines consistently showed decreased DNA damage and repair. Although XPC deficiency was associated with increased total and oxidative DNA damage, XPC knockdown was not associated with a further decrease in DNA repair or an increase in micronucleus formation or nuclear aberrancies and had little to no effect on CS toxicities including clonogenic survival and apoptosis. We hypothesize that lung cancer cells, already characterized by oncogenic mutations, may rely on alternative DNA repair mechanisms to promote survival, and evade apoptosis. Additionally, our findings indicate that lung cancer cells exhibit increased resilience to CSE compared to bronchial epithelial cells, aligning with previous studies demonstrating enhanced tumorigenicity in lung cancer cell lines following cigarette smoke exposure^29,30^.

Another novel finding is that the decrease in human XPC mRNA is not solely due to cigarette smoke exposure or field effects from lung cancer. Both lung adenocarcinoma and squamous cell carcinoma exhibit decreased XPC mRNA expression compared to non-cancerous adjacent lung tissue removed during surgery from the same subject. Although we did not investigate the underlying mechanism of decrease XPC mRNA, reductions in the 3p gene, where XPC is located, are common and occur early in lung cancer, which may account for this reduction^31^. Additionally, decreased XPC gene expression has been associated with acetylation or methylation changes at the XPC promoter site in cigarette smoke-associated bladder cancer^32,33^, although methylation changes in the promoter region were not observed in one study of non-small cell lung cancer^34^. Our findings support the hypothesis that decreased DNA repair, coupled with carcinogen exposure, contributes to lung cancer development through a double-hit mechanism, particularly in squamous cell carcinoma^35^. This suggests that reduced or erroneous DNA repair alongside exposure to cigarette smoke may lead to new mutations facilitating the transition from benign to malignant epithelial cells^7^. Collectively, these findings support a mechanism by which low XPC, decreased through exposures including cigarette smoke, plays a critical role in the early stages of epithelial cell carcinogenesis by increasing DNA damage, genomic instability, and altering the DNA damage response in these cells.

Several study limitations are important to acknowledge. *In vitro* assays utilized cigarette smoke extract, limited to aqueous tobacco smoking product constituents without *in vivo* smoke components^36^. Assay limitations for DNA damage measurement (Comet assay and CBMN) were addressed by employing multiple assays and rigorous cell counting protocols. Specific precautions were taken for each assay: an average of at least 1000 cells were counted per slide for CBMN (see Methods for more details), 10,000 cells for flow cytometry, and each experiment was replicated at least three times. To minimize variation in CSE, the cigarette smoke bubbling time was consistently maintained between at 2:00 minutes with constant flow and the same extract was used for all samples of a particular replicate and cell type. However, the impact of XPC and inhaled cigarette smoke total lung and the lung microenvironment, known to be important in lung cancer development, may differ in vivo than that observed in these in vitro studies. Further studies should confirm these findings in vivo.

## Conclusion

XPC gene expression is frequently decreased in the early stages of human lung squamous cell carcinoma and adenocarcinoma. Cigarette smoke, along with decreased NER DNA repair, plays a significant role in benign bronchial cells by increasing DNA damage and apoptosis. However, these factors have a differential effect on malignant lung cancer cells. This suggests that decreased DNA repair could play an early role in the development of the hallmark genomic instability found in lung cancers and may help explain the concurrent emphysema observed in many individuals with lung cancer.

Further translational and mechanistic studies are needed to investigate the epigenetic causes of decreased XPC gene expression in non-small cell lung cancer tumors compared to adjacent lung tissue. This could further elucidate its role in the early transition to malignant cells.

## Materials and Methods

### Chemicals and Consumables

All reagents were purchased from Thermo Fisher Scientific (Waltham, MA), unless otherwise specified. All restriction enzymes were purchased from New England Biolabs (NEB; Beverly, MA) and conducted according to the manufacturer’s instructions.

### Cell Culture

Beas-2B cells (SV40-transformed human bronchial epithelial cells; ATCC) were cultured following established protocols^12^. The non-small cell lung cancer (NSCLC) cell lines, A549 (CCL-185, Lung Carcinoma), H1299 (CRL-5803, Large Cell Carcinoma) and H520 (HTB-182, Human Lung Squamous Cancer), were procured from the American Type Culture Collection authenticated by STR testing, and cultured as previously described, incubated at 37°C in a humidified 5% CO2 atmosphere^37,38^. SV40-transformed skin fibroblast cell lines from XPC-proficient (XPC+/+, GM637), XPC-deficient (XPC−/−, GM15983), and fully corrected XPC (GM16248) individuals were obtained from Coriell Cell Biorepositories and maintained according to provided instructions^39^.

### Cigarette Smoke Extract Treatment

Cigarette Smoke Extract (CSE) treatment involved culturing cells as previously outlined^12,40^. Cells (H1299, A549, H520 at approximately 50,000 cells/cm^2^, Beas-2B at 25,000 cells/cm^2^) were seeded in coated six-well plates (9.5 cm^2^/well, Cytokinesis Block Micronucleus Assay) or standard 60mm x 15mm tissue culture dishes (21.29 cm^2^, Immunoblotting). The growth medium used was Dulbecco’s Modified Eagle Medium (DMEM)-High Glucose with 10% Fetal Bovine Serum (FBS) and 1x Streptomycin-Penicillin-Glutamine mixture. Serum starvation (DMEM-high glucose, 1x Streptomycin-Penicillin-Glutamine only) was initiated 4-5 hours after seeding. CSE preparation, involving smoke from 2 3R4F cigarettes (Tobacco Research Institute) or air (Air Control, AC) bubbled through 20mL of PBS, followed by filtration through a 0.2µm filter, was completed 16 hours after serum starvation. Cells were exposed to CSE for 24 hours.

### Transduction

The transduction procedure followed established protocols^12^. Briefly, a bronchial epithelial cell line (Beas-2B), lung carcinoma cell lines (H1299, A549), and human lung squamous cancer cells (H520) were stably transduced with lentivirus containing either shRNA (shCtrl) or shXPC RNA (A549 shXPC119A2; H1299shXPC118B1; and Beas-2B shXPC119B3) as per the manufacturer’s instructions (Sigma-Aldrich) (Figure 1). Post-transduction, cells were cryopreserved in 1x DMSO, stored in liquid nitrogen, thawed, and subsequently cultured in 75 cm^2^ cell culture flasks.

### Comet Assay

The assessment of DNA damage was conducted using the alkaline comet assay, as previously detailed^8^. To outline the procedure briefly, a 100% cigarette smoke extract (CSE) was prepared by introducing ambient air (AC) or smoke from two 3R4F cigarettes into 20 ml PBS, adjusting the pH to 7.4, and filtering through a 0.2 μm filter. Following a 16-hour period of serum starvation, cells were exposed to either AC or 5% CSE for 24 hours. The cells were then suspended in low-melting-point agarose and applied to prepared coverslips. After solidification, the cells were lysed, and DNA unwinding occurred using an alkaline solution (2.5 M NaCl, 100 mM ethylenediaminetetraacetic acid, 10 mM Tris, 1% Triton X-100, pH 10), followed by single-cell electrophoresis, in accordance with the manufacturer’s instructions (Trevigen). Slides were dehydrated using alcohol and stained with SYBR gold. Images were captured at 10x magnification (fluorescein isothiocyanate filter) using a Nikon Eclipse 90i microscope, and digital photographs were obtained using NIS Elements.

Analysis of a minimum of 50 comets was conducted using CometScore (TriTek). The mean tail DNA percentage from three independent experiments was compared using analysis of variance.

The assessment of base excision repair (BER) OGG1 activity was carried out through the Alkaline Comet FLARE assay (Trevigen), both immediately after and 24 hours following incubation with cigarette smoke extract^41^.

### Cytokinesis-Block Micronucleus Assay (CBMN)

The CBMN protocol was adapted from Farabaugh, Doak, Roy, and Elespuru^42^. After CSE or AC (filtered air) exposure, cells were detached using Trypsin-EDTA (0.25%) followed by three successive rounds of centrifugation (200g, 10min), with resuspension in 1mL Phosphate Buffered Saline (PBS) after each centrifugation. Upon completion, cells were seeded on six well plates with fibronectin-coated glass coverslips (Corning® Fibronectin, human, 5mg), and 3.5µg/mL Cytochalasin-B (CAS14930-96-2) was added to each well with media. After 24 hours, media was removed, and cells were incubated with 900µL PBS, 900µL 0.075KCl, and 200µL methanol/glacial acetic acid 25:1 for 10 minutes, followed by incubation with 1mL of the methanol/glacial acetic acid 25:1 mixture for 10 minutes. Coverslips were mounted on glass slides with Permanent Mounting Medium containing DAPI. Cells were visualized and imaged via fluorescent microscopy for DAPI immunofluorescence.

### Cell Counting and Image Analysis

Images were taken and total cell count for each image was determined using ImageJ Batch Processing. Cells on the picture border were excluded. Outliers were removed using the Remove Outliers tool on ImageJ. Images were manually evaluated for micronuclei and nuclear blebs. The parameters for micronucleus scoring established by Fenech^43^ were used to quantify micronuclei: 1) micronuclei are no larger than 1/3 the size of the true nucleus, 2) micronuclei are distinguishable from tissue artifact in that they are non-refractile, 3) micronuclei are discrete entities that should not be attached to the main nucleus, 4) the boundary of the micronucleus must be clearly distinct from that of the main nucleus, and 5) micronuclei should stain as intensely, if not more intensely, than the main nucleus. Nuclear blebs were defined as distinct, deforming out pockets of the nucleus that protrude significantly and cannot be accounted for simply by abnormalities in nuclear shape. Total nuclear aberrations were calculated as the combined total of cells with micronuclei and nuclear blebs. To offset the impact of bias and human error, the average number of cells counted per slide was 1156.4, 1744.2, and 1413.2 for Beas-2B, H1299, and A549, respectively.

### Protein Extraction and Immunoblotting

Immunoblotting was conducted on whole cell extracts following established procedures^37^, employing validated antibodies. Densitometry analysis was performed using Image Lab Software from Bio-Rad.

### Plasmids

Plasmids used in these experiments have been previously optimized for HCR assays in our laboratory^16^. We used pMAX-GFP (Lonza, Cologne, Germany), that encodes for a green fluorescent protein and pCMV-E2-Crimson (Takara Bio, CA, USA), which encodes for E2-Crimson (far red).

### Transfection

All transfections were done via nucleofection using 4D-Nucleofector X-Unit (Lonza, Cologne, Germany). Transfections were done according to the manufacture protocol with modifications. Transfection was optimized for each cell line in 20 µL 16-well strips using SF solution/EH-100 program with 0.2 µg of each plasmid for 0.3-0.5×10^6^ cells used for each transfection.

### Flow Cytometry

All flow cytometry was performed on a FACSCalibur flow cytometer (BD Biosciences, USA). Cells were visualized on a FSC vs SSC dot plot after that a gate was placed around the desired population excluding dead cells and debris and at least 10,000 gated cells were counted. Controls for each experiment included single plasmid and mock-transfected cells to determine transfection efficiency and confirm positive and negative fluorescence expression for gating by flow.

### Measurement of DNA Repair through Host-Cell Reactivation Assay

Assay was performed according previously optimized protocol^16^. For determination of NER repair, a plasmid producing green fluorescence (pMAX-GFP) was mock-treated (control) or modified by treatment with 10 J/m2/s of ultraviolet-C (UV) through a germicidal lamp^12^. NER repair was measured by flow-cytometry after transfection with a modified or unmodified plasmid (pMAX-GFP) and co-transfection with a second unmodified, covalently closed plasmid producing “far red” fluorescence to control for transfection efficacy (pCMV-E2-Crimson). The relative GFP expression is determined by dividing all green cells (green alone + [green and red]) by all red cells (red alone + [green and red]) [the “relative GFP expression”]. The NER% (percent efficiency) was determined by dividing relative GFP expression in cells transfected with the modified GFP plasmid by the relative GFP expression in cells transfected with the unmodified (control) GFP plasmid.

### Human Gene Expression Analyses

XPC mRNA expression in unmatched non-small cell lung cancers filtered by histologic subtype (adenocarcinoma or squamous cell carcinoma) or “normal” (non-cancerous) lung, and associated clinical data, were downloaded from publicly available datasets deposited in GEO and TCGA (GEO ID/Source TCGA_LUAD and TCGA_LUSC) and analyzed using Lung Cancer Explorer^44^.

Frozen human tissue samples comprised of previously untreated lung adenocarcinoma or lung squamous cell carcinoma, and subject-matched non-cancerous lung removed at the time of surgery were obtained from the Indiana University Simon Comprehensive Cancer Center (IUSCCC) Tissue Bank through an IRB 1709430417). Samples were deidentified with available demographic data available for all subjects unless otherwise noted, including subject age, ethnicity, race, gender, and cigarette use (current, former, or never). All specimens were reviewed by a board-certified pathologist for histology (tumor type and no evidence of malignancy) and suitability for analysis. RNA was extracted using TRIzol and mechanical homogenization, then purified by QIAGEN RNeasy Mini Kit. Spectrophotometric analysis of RNA quality and concentration preceded reverse transcription to cDNA. Relative quantification in real-time polymerase chain reaction (qRT-PCR) was performed with TaqMan Gene Expression Master Mix and primers (Applied Biosystems) for XPC (Hs01104206_m1) and an endogenous control (GAPDH, 4333764F) as published^12^. Matched samples were run on the same plate with an inter-plate control (BEAS-2B cDNA), each performed in triplicate.

### Data Processing and Statistical Analysis

At least three replicates of each experiment, for each cell type, were completed. One-way and two-way Analysis of Variance (ANOVA) were completed using SigmaPlot v14.5. Pairwise Multiple Comparisons were completed via the Holm-Sidak method on SigmaPlot. Statistical significance was considered as p<0.05. Figures were generated using CoreIDRAW X6.

## Abbreviations

8-oxoG: 8-oxo-7,8-dihydro-2’-deoxyguanosine
ANOVA: Analysis of Variance
BER: base excision repair
CBMN: cytokinesis-block micronucleus
cDNA: complementary DNA
CMV: cytomegalovirus
COPD: chronic obstructive pulmonary disease
CS: cigarette smoke
CSE: cigarette smoke extract
DNA: deoxyribonucleic acid
FLARE: Fragment Length Analysis using Repair
FSC: forward scatter
GEO: Gene Expression Omnibus
GFP: green fluorescence protein
hOgg1: human 8-oxoguanine DNA glycosylase
IRB: Institutional Review Board
IUSCCC: Indiana University Simon Comprehensive Cancer Center
LUAD: lung adenocarcinoma
LUSC: lung squamous cell carcinoma
mRNA: messenger RNA
NER: nucleotide excision repair
NSCLC: non-small cell lung cancer
NTCU: N-nitroso-tris-chloroethylurea
PARP: poly (ADP-ribose) polymerase
PI: propidium iodide
RT-qPCR: quantitative reverse transcription polymerase chain reaction
SSC: side scatter
shCtrl: cell lines stably expressing non-targeted, scramble control short hairpin RNAs
shXPC: stable knock-down of XPC using short hairpin RNA
TCGA: The Cancer Genomic Atlas
UV: ultraviolet
XPC: Xeroderma Pigmentosum Group C

## Author Contributions

Conceptualization, C.R.S.; data curation, N.A.N., R.S.S. and C.R.S.; formal analysis, N.A.N., R.S.S. and C.R.S.; funding acquisition, C.R.S.; investigation, N.A.N., B.L., B.M.W., M.N.K., E.A.M., H.Z., N.P.; methodology, C.R.S., M.W.G and R.S.S.; project administration, C.R.S.; resources, C.R.S. and M.W.G.; software, R.S.S.; supervision, C.R.S. and R.S.S.; validation, N.A.N., R.S.S. and C.R.S.; visualization, N.A.N., B.L., B.M.W., M.N.K. and C.R.S.; writing—original draft, N.A.N., B.L., B.M.W., C.R.S.; writing—review and editing, N.A.N., B.L., B.M.W., M.N.K., E.A.M., H.Z., N.P., R.S.S., M.W.G.; All authors have read and agreed to the published version of the manuscript.

## Acknowledgments

We appreciate the Department of Medicine, Indiana University School of Medicine, who provided dedicated research time to B.L. during her Internal Medicine Residency Training.

## Ethical Statement

The study was approved by the Institutional Biosafety Committee of Indiana University School of Medicine (protocol #IN-972) and the Richard L. Roudebush Veterans Affairs Medical Center (protocol #21018). No human experimentation was performed in this study. Frozen human samples were obtained from the Indiana University Simon Comprehensive Cancer Center (IUSCCC) Tissue Bank through an IRB (#1709430417). No AI was used in the writing or production of this manuscript.

