## Supplemental Figures and Tables for "Cigarette Smoke and Decreased DNA Repair by Xeroderma Pigmentosum Group C Use a Double Hit Mechanism for Epithelial Cell Lung Carcinogenesis"

### Supplemental Figures and Methods

#### Supplemental Figures:

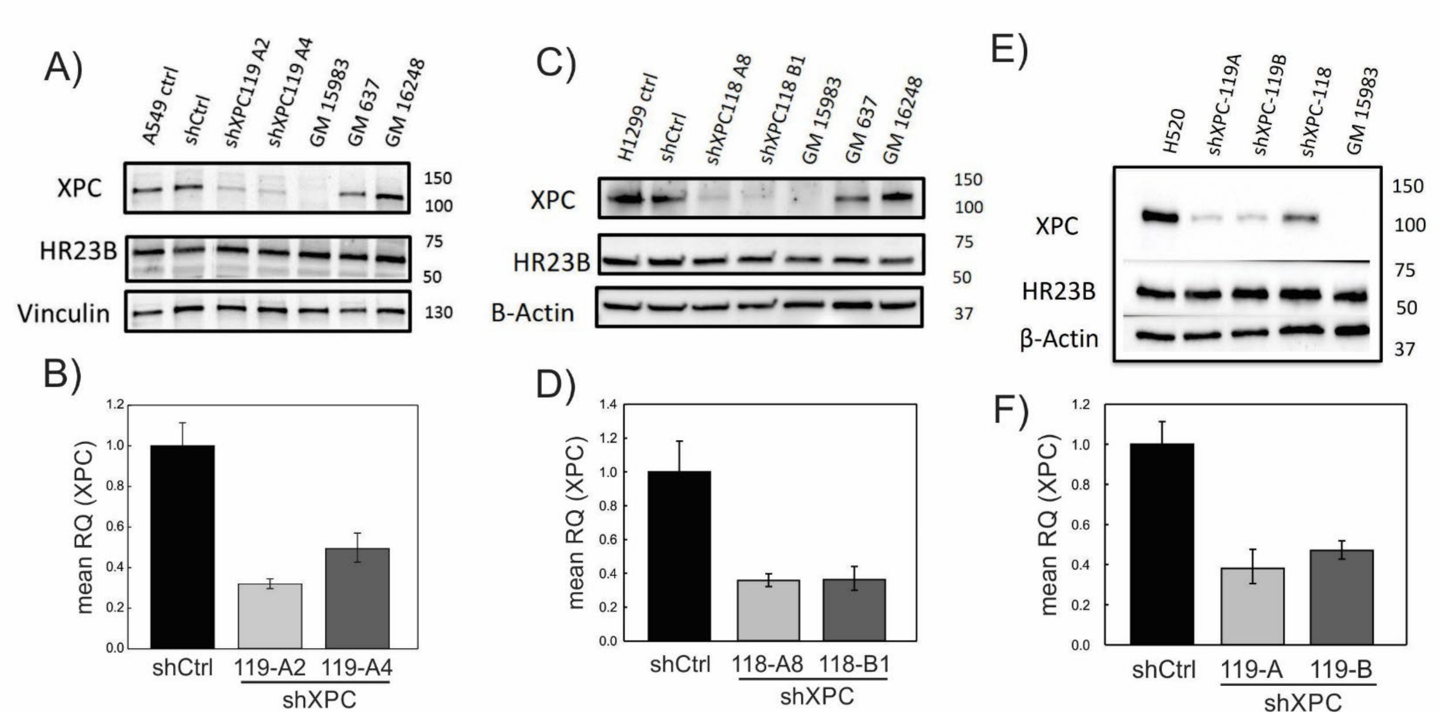


**Supplemental Figure 1:** Stable XPC knock-down cell line development. Lentiviral siRNA to XPC performed in 3 cancer cell lines: A549 (A,B), H1299 (C,D) and H520 (E, F). Western blot analysis shows decreased XPC protein (A, C, E) expression. Decreased *XPC* mRNA expression by RT-qPCR (B, D, F) expression. shCtrl (siRNA to non-targeted XPC expression levels are measured by Western blot analysis in A549 cells after stable KD of XPC by lentiviral shRNA and compared with loading control (β-actin), with specificity confirmed by expression levels of the XPC complexing protein, HR23B (A). Two XPC KD clones (A549: shXPC119-A2 and 119-A4 are shown with the parent cell line (A549), nontargeting shRNA control (shCtrl) and control fibroblast cell lines from an unaffected human (GM637), and XPC-deficient proband unmodified (GM15983) and reconstituted with XPC cDNA (GM16248). Calculated XPC expression levels normalized to loading control and compared with shCtrl are shown below a representative immunoblot. XPC RNA expression levels by qRT-PCR are shown compared with shCtrl (B). Likewise, (C) showed XPC expression levels as measured by Western blot analysis in H1299 cells as well as two XPC KD (shXPC 118 A8 and shXPC 118 B1). (D) shows the decreased XPC RNA expression normalized to shCtrl. Finally, (E) showed XPC expression levels as measured by Western blot analysis in H520 cells as well as two XPC KD (shXPC 119A and shXPC 119B) and (F) shows the decreased XPC RNA expression normalized to shCtrl. (mean ± SD; n = 3). MW = molecular weight.

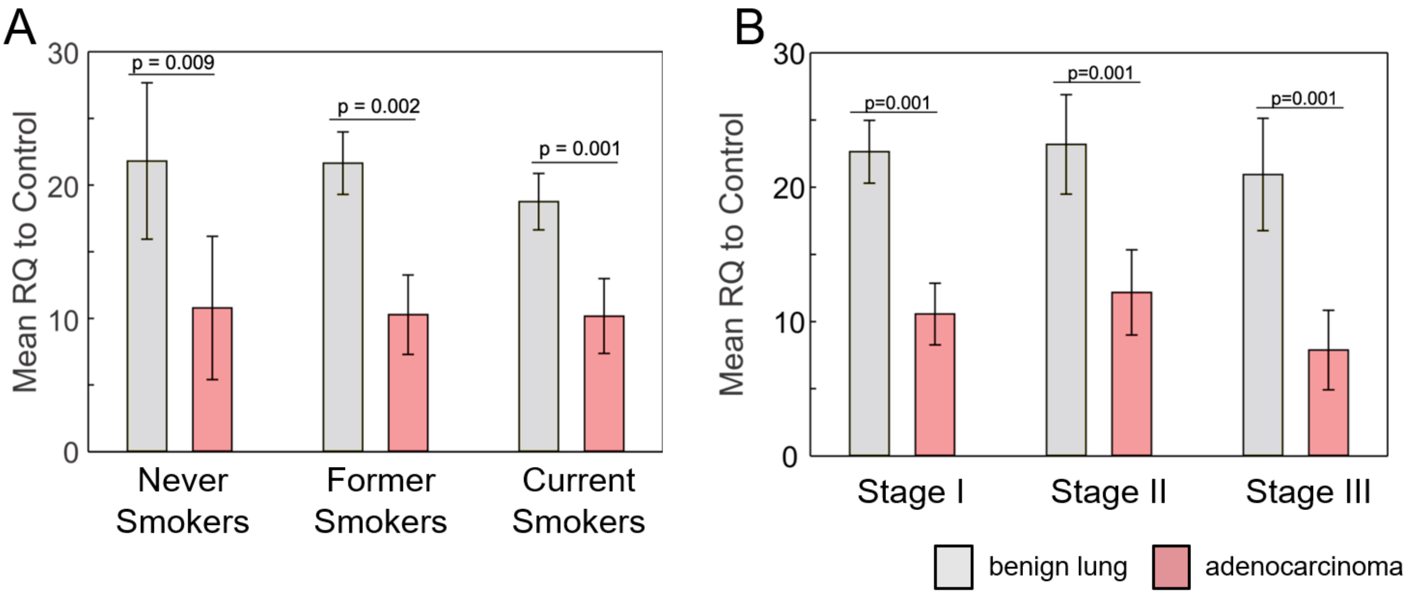


**Supplemental Figure 2:** Subgroup analysis of XPC mRNA expression divided by A) smoking status and B) stage.

#### Supplemental Tables:

**Table 1: Human lung adenocarcinoma samples - Demographics**

| Demographics | Percent/Average (SD) |
|---|---|
| Number of subjects (matched specimens) | 45 (44 used for analysis)* |
| Age | 63.5 (+/- 10.8) |
| Male Sex | 43% |
| Race | |
| White | 82% |
| Black | 16% |
| Asian | 2% |
| Cigarette smoking status | |
| Ever Smoker | 61.4% |
| Never Smoker | 20.5% |
| Current Smoker | 16% |
| Unknown/Not recorded | 22.7% |
| Stage at time of tissue collection | |
| Stage I | 41% |
| Stage II | 39% |
| Stage III | 14% |
| Stage IV | 2% |
| Unknown/Contradictory | 5% |

*insufficient/poor quality RNA yield from tumor

**Table 2: Human lung squamous cell carcinoma samples - Demographics**

| Demographics | Percent/Average (SD) |
|---|---|
| Number of subjects (matched specimens) | 36 |
| Age | 63.9 (+/- 7.7) |
| Male Sex | 56% |
| Race | |
| White | 94% |
| Black | 6% |
| Cigarette smoking status | |
| Ever Smoker | 86.1% |
| Never Smoker | 0% |
| Current Smoker | 41.7% |
| Unknown/Not recorded | 13.9% |
| Stage at time of tissue collection | |
| Stage I | 33% |
| Stage II | 42% |
| Stage III | 17% |
| Stage IV | 0% |
| Unknown/Contradictory | 8% |
